# Flexible Modulation of Neural Variance Facilitates Neuroprosthetic Skill Learning

**DOI:** 10.1101/817346

**Authors:** Albert K. You, Bing Liu, Abhimanyu Singhal, Suraj Gowda, Helene Moorman, Amy Orsborn, Karunesh Ganguly, Jose M. Carmena

## Abstract

One hallmark of natural motor control is the brain’s ability to adapt to perturbations ranging from temporary visual-motor rotations to paresis caused by stroke. These adaptations require modifications of learned neural patterns that can span the time-course of minutes to months. Previous work with brain-machine interfaces (BMI) has shown that over learning, neurons consolidate firing activity onto low-dimensional neural subspaces, and additional studies have shown that neurons require longer timescales to adapt to task perturbations that require neural activity outside of these subspaces. However, it is unclear how the motor cortex adapts alongside task changes that do not require modifications of the existing neural subspace over learning. To answer this question, five nonhuman primates were used in three BMI experiments, which allowed us to track how specific populations of neurons changed firing patterns as task performance improved. In each experiment, neural activity was transformed into cursor kinematics using decoding algorithms that were periodically readapted based on natural arm movements or visual feedback. We found that decoder changes caused neurons to increase exploratory-like patterns on within-day timescales without hindering previously consolidated patterns regardless of task performance. The flexible modulation of these exploratory patterns in contrast to relatively stable consolidated activity suggests a simultaneous exploration-exploitation strategy that adapts existing neural patterns during learning.

## INTRODUCTION

Our brains have an intrinsic ability to adapt to perturbations. However, the degree to which we can adapt depends on the similarity to previously learned behaviors and the time allowed for adaptation (Izawa et al., 2008; Kording et al., 2007; Shadmehr et al., 2010; Wei and Körding, 2009). Brain-machine interfaces (BMI) allow us to interrogate specific populations of neurons and examine how they adapt during learning (Athalye et al., 2020; Golub et al., 2016; Shenoy and Carmena, 2014). BMI decoders translate neural activity ranging from firing rates to field potentials into actions taken by an end effector (e.g. computer cursor or robotic arm). It has been shown in multiple studies that the brain adapts and forms stereotyped firing patterns over learning (Carmena et al., 2003; Chase et al., 2012; Ganguly and Carmena, 2009; Jarosiewicz et al., 2008; Koralek et al., 2012; Orsborn et al., 2014; Taylor, 2002; Wander et al., 2013). We refer to these stereotyped patterns as neuroprosthetic maps (Ganguly and Carmena, 2009; Ganguly et al., 2011; Orsborn et al., 2014). More recent work has shown how the brain coordinates the activity of neurons over learning and collapse upon a low-dimensional neural subspace as performance improves when given a fixed decoder (Athalye et al., 2017; Oby et al., 2019).

In addition, similar results regarding neuroprosthetic map formation have been shown in BMI experiments where decoders are periodically readapted to fit changes in the neural population (Orsborn et al., 2014). We refer to these paradigms as “2-learner systems.” In these experiments, closed-loop decoder adaptation (CLDA) was used to refit decoder weights to updated neural intentions at the start of the day, allowing for quick improvements in BMI performance (Dangi et al., 2013; Gilja et al., 2012; Moorman et al., 2017; Orsborn et al., 2012, 2014). However, while CLDA incrementally fits the overall control space closer to the neural subspace, the decoder weights of individual neurons could be affected due to changes in the ensemble of neurons, the variance of neural activity, or sources of noise. Thus, new decoder weights inherently introduce slight perturbations to individual neurons even if the overall population activity is better modeled. In these experiments, task performance continued to improve alongside these decoder refits. Thus, we asked how neurons adapt given slight decoder changes throughout learning. Furthermore, what is the timescale of this adaptation? Refitting a new decoder happens on the timescale of minutes. Thereafter, the decoder is fixed for the duration of the day and sometimes held fixed for multiple days. How do neurons respond to relatively fast changes in decoder weights and still maintain the ability to consolidate firing patterns over longer multi-day timescales?

Since CLDA fits to the neural intentions of the subject, one can assume that any changes in weights in the decoder are mainly rotations within the low-dimensional neural subspace, sometimes referred to as the intrinsic manifold. As reported in recent studies, such perturbations are quickly learned within-day whereas out-of-manifold perturbations are more difficult for neurons to adapt indicating resistance to changes in the neural subspace (Golub et al., 2018; Oby et al., 2019; Sadtler et al., 2014). Thus, neurons may be adapting to periodically refit decoders as they would to fixed decoders since the control spaces are not significantly perturbed by new decoder weights. However, these decoder weight changes introduce errors on individual neurons such as misalignment of preferred tuning directions, potentially creating slight out-of-manifold perturbations. In order to adapt to these errors, we hypothesize that neurons increase their neural firing rate variance while maintaining relatively stable, coordinated population-level firing patterns on short, within-day timescales.

To observe the effects of perturbations on the consolidation of neural firing patterns throughout learning, we analyzed data from three BMI experiments performed on a total of five male rhesus macaques. These experiments refit decoders at various frequencies, ranging from repeated decoder swaps within-day to performing CLDA at the beginning of each day (Figure 1). These differences allowed us to gain increased insight on how neurons adapt on short versus long-term timescales. As expected, our results show that neurons increased neural variance immediately after decoder-refitting. However, the variance dropped as the decoder was held fixed, thereby increasing the fraction of neural variance that was aligned with the low-dimensional manifold. Importantly, these manifolds did not change within day nor across many days. Rather, neural variance was slowly collapsed onto a relatively stable neural subspace over learning. These flexible changes in neural variance alongside stable low-dimensional neural spaces from the onset of learning suggest a concomitant exploration and exploitation strategy during motor learning, where new neural patterns are formed by adapting preexisting patterns.

**Figure 1:**
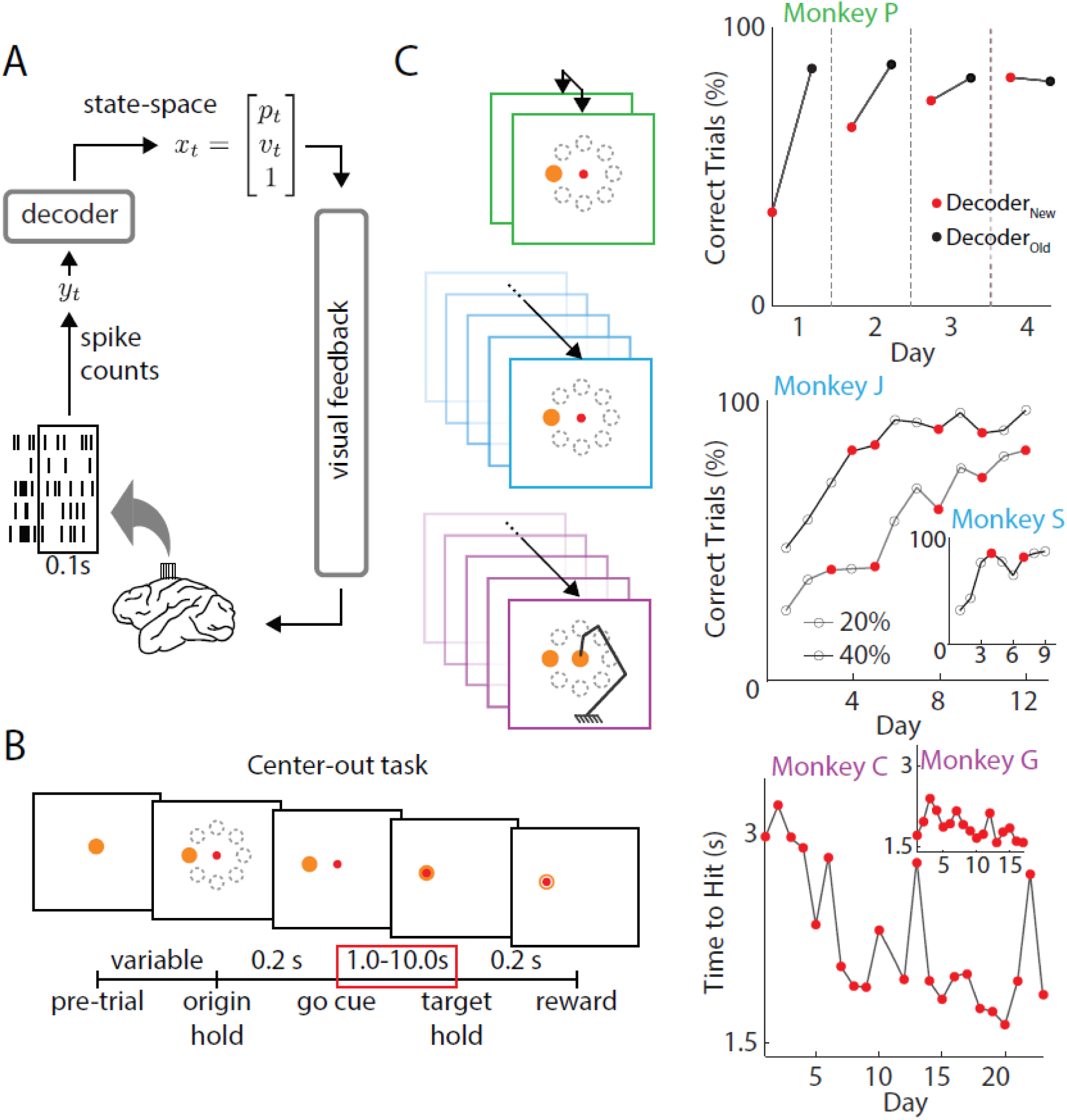
Task design and performance. A. Closed-loop BMI. Neural signals were recorded from motor cortices in the brain of rhesus macaques. Spikes were recorded and decoded to update a state-space vector, moving the effector in the animals’ visual field (adapted from Moorman and Gowda et al., 2017). B. In all three experiments, monkeys were instructed to perform a center-out task, moving the effector from a center target to one of eight randomly selected peripheral targets. Successful trials were rewarded with juice. Only rewarded trials during movement periods (red box) were analyzed in this study (adapted from Moorman and Gowda et al., 2017). C. Three experiments were analyzed in this study. *Green*, two decoders were used each day (Ganguly and Carmena, 2009). After proficient control of one decoder (black), a new decoder was introduced at the beginning of each successive day (red). After a period each day, decoders were swapped back to the previously learned decoder. Percent accuracies were calculated as the ratio of successful trials to number of self-initiated trials. *Blue*, decoders were re-adapted periodically across training and neural populations occasionally changed from day-to-day (Orsborn et al., 2014). Monkeys were trained to perform a center-out task with decoders of varying amounts of adaptation (CLDA) as defined by the initial performance of each time series (i.e. 20% and 40%). Monkey J was trained on two time series. After 12 days of 20% CLDA, he was switched to a 40% decoder. Monkey S remained on a 20% decoder for the entirety of the experiment. Red dots indicate decoder re-adaptation. *Purple*, a virtual arm with 4 degrees of freedom (DOF) was controlled with a new decoder each day (Moorman and Gowda et al., 2016). Monkeys trained to move the effector in 2D space and improved reach times over learning (Monkey C, R^2^ = 0.431, p = 6.6e-4; Monkey G, R^2^ = 0.298, p = 0.0233). CLDA was performed each day and percent accuracy was saturated from Day 1 for both animals (∼80% correct trials across all days).

## RESULTS

In this study we analyzed data from three previous studies (Ganguly and Carmena, 2009; Orsborn et al., 2014; Moorman and Gowda et al., 2017) involving five monkey subjects. In each experiment, decoders were periodically changed and monkeys were trained over the time-course of days thus allowing us to ask how neural consolidation occurs alongside changes in decoder weights over learning. Furthermore, all three experiments consisted of center-out tasks in 2D space whereby monkeys were instructed to move the effectors to eight peripheral targets (Figure 1B). In Experiment 1 (Figure 1C, green), two biomimetic decoders were used each day. The animal (Monkey P) had previous experience with one decoder (Decoder_OLD_) but was naïve to the second (Decoder_NEW_). While performance with the old decoder was stable and high across all four days, control with the new decoder improved across days (Figure 1C, green).

In Experiment 2 (Figure 1C, blue), both monkeys (J and S) learned to control the BMI with suboptimal decoders as indicated by their initial performance on Day 1 (Figure 2B). Closed-loop decoder adaptation (CLDA) was used to intermittently refit decoder weights over the time-course of learning (Figure 1C, middle, red dots). Note that while Monkey J had two time series of data collected, the 40% CLDA condition (indicating 40% performance accuracy after CLDA training on the first day) occurred the day after the 20% CLDA condition was completed.

**Figure 2:**
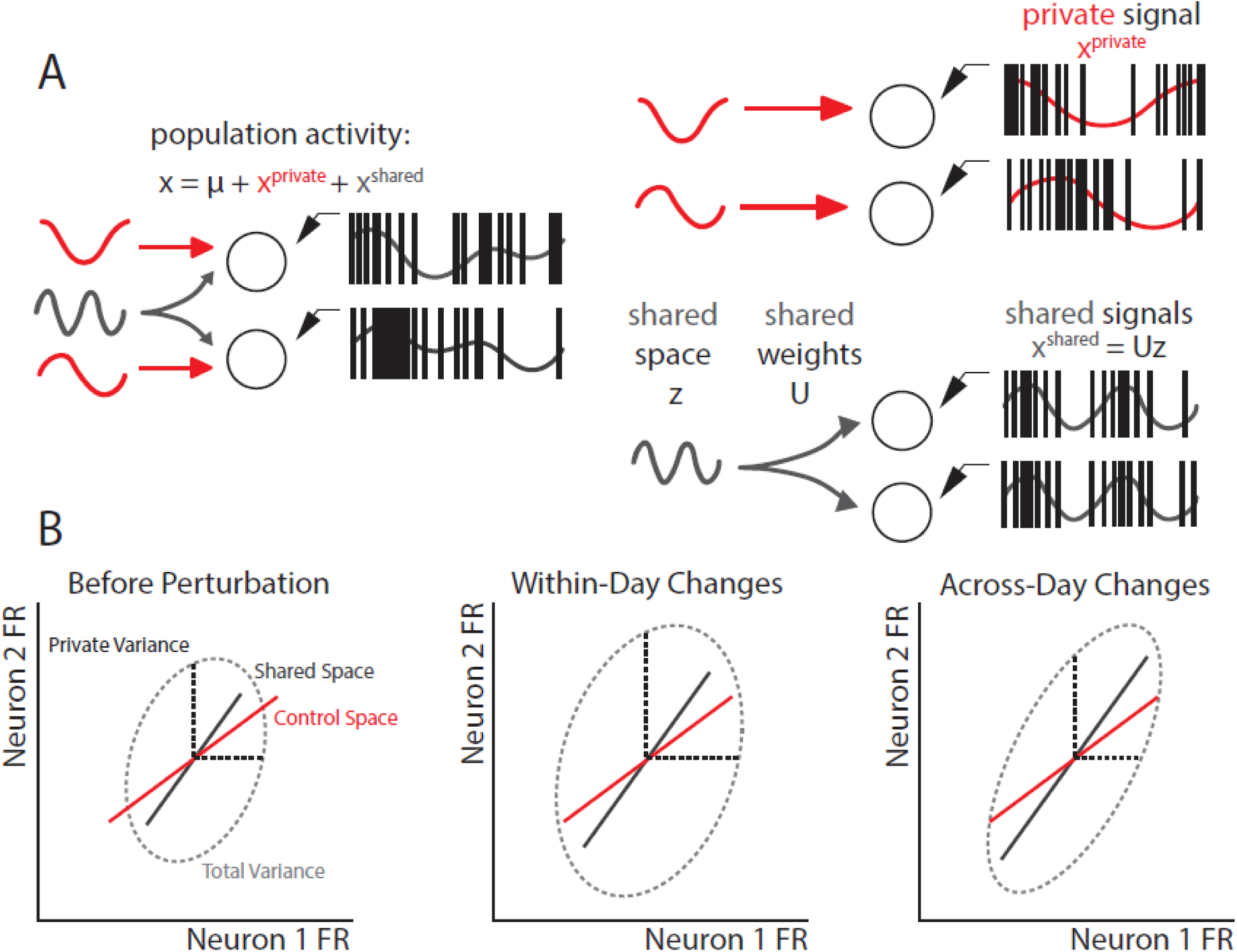
Factor analysis model predictions. A. Factor Analysis (FA) model. Neural signals can be demeaned (*μ*) and decomposed into covarying signals (shared) and uncorrelated signals (private). Intuitively, shared signals indicate a low-dimensional neural space where neurons fire in coordinated ways (Athalye et al., 2017). B. Predictions of neural subspace changes in response to decoder re-adaptation. Neural firing patterns between two neurons are shown in the dashed ellipses. Private variance (black, dashed) explains the variance that is independent from other neurons. Before decoder re-adaptation, there is an intrinsic shared space (gray). Since CLDA fits to neural intentions, we expect the decoder plane (control space, red) to be fairly aligned with the neural subspace (shared space, gray). Lengths of lines indicate amount of variance captured by each component. Immediately following decoder re-adaptation, we expect no changes in the angle of the shared space. Rather, we expect an increase in the overall neural variance in order to better capture the control space. Note that this increases the shared variance, but not the ratio of shared to total variance. Over multiple days of learning with the same (or similar) decoder, we expect consolidation patterns to occur with increased shared activity (longer gray line) and decreased private activity.

Finally, in Experiment 3 (Figure 1C, purple), two monkeys (C and G) moved a kinematically redundant 4 degree-of freedom (DOF) virtual arm to the same set of targets as in the previous experiments. CLDA was performed for roughly 10 minutes at the beginning of each day leading to high performance from the first day. However, decoders weights were still changed from day to day; correlations of these weights are shown in Supplemental Figures 4,5. While the percent correct was saturated from the first day (∼80% for both animals), their reach times continued to decrease across training indicating skill improvement (Figure 1C, purple).

### Factor Analysis Predicts Flexible Modulation of Consolidated Patterns

As it has been previously reported (Athalye et al., 2017; Golub et al., 2018; Oby et al., 2019; Sadtler et al., 2014), factor analysis (FA) provides an intuitive model for how neurons coordinate firing patterns. In short, FA analyzes population-level firing trends and decomposes each neuron’s firing activity into covarying (shared) and uncorrelated (private) components (Figure 2A). Importantly, since we were interested in how neural populations adapted over learning, only units that were stably recorded across the entire timeseries for each experiment were used. Geometrically, this shared space (or often referred to as an intrinsic manifold) refers to a low-dimensional neural subspace where neurons fire in correlated ways (Figure 2B). Subsequently, the private variance (PV) captures the variance of firing rates of a neuron that is independent from other neurons and has been suggested to be a correlate of exploratory patterns (Athalye et al., 2017).

Previous work in the field has focused primarily on the effects of BMI learning on shared spaces and it has been shown that decoders that align with the neural shared space (i.e. intrinsic manifold) can be learned more readily within a day whereas decoders that are explicitly out of alignment with the intrinsic manifold cannot be as quickly learned (Sadtler et al., 2014). Since our experiments rely on biomimetic movements and CLDA to fit effector kinematics to the neural intentions of the animal, we would expect the decoders used in these experiments to be roughly within the intrinsic manifold and therefore we would expect stable alignment of the shared space (Figure 2B). Hypothetically, if a decoder was perfectly fit to the neural intentions of an animal, we would expect no changes in either the angle or the size of the shared space. However, due to the high dimensionality of the neural populations recorded during these BMI tasks, decoder weights change slightly upon a decoder refit. Thus, we hypothesize that since the intrinsic manifold is relatively stable on short timescales, neurons must increase their overall firing rate variance (including both PV and shared variance) to accommodate perturbations in the control space (i.e. decoder weights) (Figure 2B, middle). Subsequently, decoders that are less aligned with the intrinsic manifold would cause a larger increase in overall neural variance.

Additionally, work by Athalye et al., 2017 showed that neurons increased their proportion of shared variance to total variance (SOT) over learning with a fixed decoder, indicating an emergence of coordinated or consolidated neural activity. While CLDA is used to periodically adapt the decoder weights, animals are able to learn the task over the time course of days (Orsborn et al., 2014). Together, this leads to the hypothesis that SOT increases even in the context of a 2-learner system (Figure 2B, right). Furthermore, since task performance is comparable before and after CLDA, some neural adaptation (e.g. changes in total variance) must occur within the same day (Figure 2B, middle). In summary, we expect the neural variance of BMI input neurons to increase as a function of decoder unfamiliarity on a short timescale while shared structures are preserved and consolidated over a longer timescale.

### Neural Variance Flexibly Changes

Past work has suggested that PV is linked with exploratory neural patterns that are less goal-potent than shared activity but sufficient for achieving target hits (Athalye et al., 2017). Since decoder refitting inherently causes some small perturbations in the decoder readout, we would expect exploratory patterns to be engaged to accommodate increases in error. Congruent with past results, we found that the PV increased each time the monkeys controlled the BMI with a new decoder (Figures 3A,C,E). In Experiment 1, training with DecoderNEW led to higher PV in each of the four days compared to training with DecoderOLD, which was already well-learned (t-test, p = 1.3e-7). In Experiment 2, the PV was calculated each day the decoder was refit and all days following the perturbation (Figure 3C). The PV consistently increased the day following a decoder refit for both monkeys, though only Monkey J had enough perturbations to test for significance (t-test; Monkey J, p = 0.0334). Lastly, in Experiment 3, where a new decoder was refit each day, the magnitude of PV change was correlated with the amount of change in decoder weights (Figure 3E). Decoders that were similar to ones previously learned resulted in smaller PV changes (correlation; Monkey C, R = −0.70, p = 2e-4; Monkey G, R = −0.63, p = 6.3e-3). Implicit in this result is that PV decreases when more familiar (or fixed) decoders are used. We found that indeed PV decreased when the task switched to using decoders that were more familiar (Supplemental Figure 1A) or fixed for a period of days (Supplemental Figure 1B) echoing results from Athalye et al., 2017.

**Figure 3:**
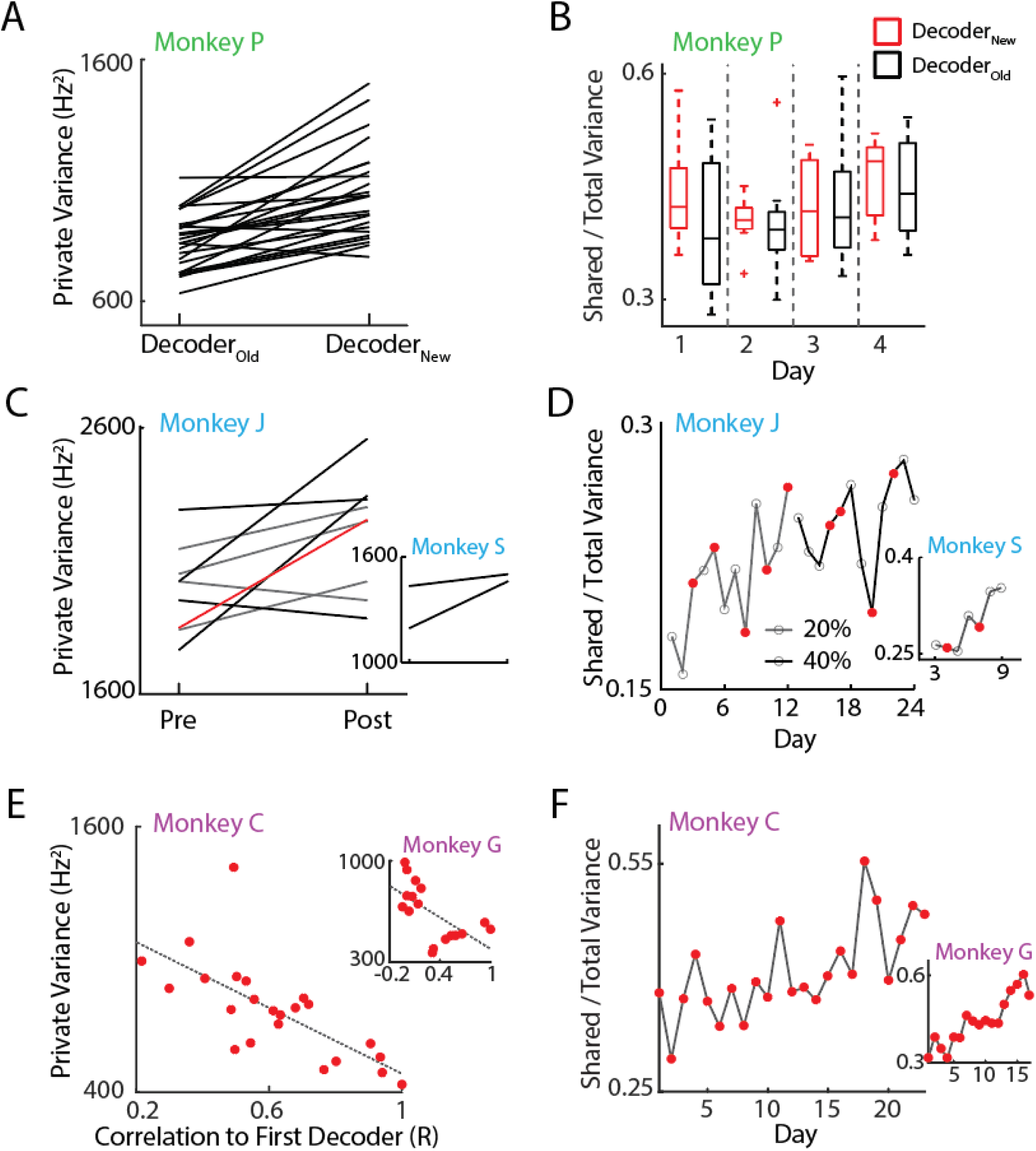
Exploratory and consolidated patterns. A. Private variance (PV) changed within day. Using the new decoder increased the PV at the beginning of each day (t-test; p = 1.5e-6). B. Consolidated patterns did not change within day. The shared-to-total variance ratio (SOT) was stable each day and did not change as a result of switching decoders. C. PV increases when decoders are re-adapted. Across both 20% and 40% time series, PV increased when decoders were refit with CLDA. The red line indicates the transition from the 20% experiment to the 40% experiment. (Wilcoxon rank-sum test; Monkey J, p = 0.0315; Monkey S n.s., only two decoder re-adaptations were performed). D. SOT increased over both time series across days (linear regression; Monkey J, R^2^ = 0.46, p = 2.8e-4; Monkey S, R = 0.81, p = 6.1e-3). E. PV was higher in cases where decoder weights were more different from the first day. (correlation; Monkey C, R = −0.70, p = 2e-4; Monkey G, R = −0.63, p = 6.3e-3). F. SOT increased over days with re-adapted decoders each day. (linear regression; Monkey C, R^2^ = 0.435, p = 3e-4; Monkey G, R^2^ = 0.829, p = 3.9e-7).

Remarkably, these changes in PV were exhibited on the timescale of minutes. Private variance decreased within the same day when an unfamiliar decoder was replaced with a well-learned decoder (Supplemental Figure 1A). Conversely, switching back to the unfamiliar decoder the following day increased the PV (Figure 3A). These results show the flexibility of neural variance, which may be required when adapting to new tasks or decoders.

### Neural Exploration Adapts Consolidated Patterns

We asked how neural consolidation emerged in a 2-learner system despite frequent changes in neural firing patterns due to changes in decoder weights. To track the proportion of neural activity that fired in coordinated patterns, we normalized the shared variance by the total neural variance (SOT). Thus, increases in SOT act as a measure for how covaried neurons’ firing activities are and can be thought of as the degree of consolidation of neural patterns (Athalye et al., 2017). We found that while PV was modulated due to decoder changes, the SOT remained stable on within-day timescales in Experiment 1 where decoders were switched in the middle of the day (Figure 3B). Together, these results indicate a scaling of the total variance (including shared variance) when a perturbation arises as predicted in Figure 2B (middle). In addition, SOT increased in experiments where a single decoder was used each day (Figures 3D, F) (regression, Monkey J, R^2^ = 0.46, p = 2.8e-4; Monkey S, R^2^ = 0.81, p = 6.1e-3; Monkey C, R^2^ = 0.435, p = 3e-4; Monkey G, R^2^ = 0.829, p = 3.9e-7). The decoupling of changes in PV and SOT over different timescales suggests that neural exploration and exploitation can simultaneously occur, and increased exploration does not occur at the expense of consolidated patterns. In contrast, increases in SOT over the time-course of many days despite flexible changes in PV may suggest that exploratory patterns consolidate over time and that neurons are adapting learned behaviors rather than generating completely novel patterns.

Furthermore, we asked if the shared spaces were stable across different decoders or if poor performance with a new decoder would lead to adaptation to a different low-dimensional space. To answer this question, we computed the “shared alignment” between the neural subspaces used to control the two decoders in Experiment 1 (Supplemental Figure 2A). We found that the shared spaces used to control the task were nearly identical despite large differences in task performance (Supplemental Figure 2B). Similarly, we compared the shared alignment between adjacent days and across days for Experiments 2 and 3 and found that shared spaces were stable on short and long timescales (Supplemental Figure 2B-D). In all cases, the shared alignment was high in adjacent blocks or days (∼0.8 across all animals), suggesting that neurons rely heavily on preexisting patterns regardless of task performance.

### Modulation Depth Facilitates Changes in Neural Variance

Finally, we asked how neurons are able to flexibly modulate their neural variance on short, within-day timescales. Intuitively, our results imply that the dynamic range of firing rates across all neurons must also be flexible to account for changes in neural variance. That is, solely increasing the mean firing rate of neurons does not necessarily increase the variance. Rather, we would expect some modulation of firing rates. Past literature has shown preferred direction tuning in BMI neurons. The dynamic range of a neuron’s tuning curve is often referred to as the modulation depth (MD) of the neuron (Carmena et al., 2003; Ganguly and Carmena, 2009; Ganguly et al., 2011; Orsborn et al., 2014). Past work has shown MD to change over many days of BMI learning. However, given the flexibility of the neural variance, we hypothesized that MD could also change on a shorter timescale.

We calculated the preferred tuning models of each neuron by fitting a cosine tuning curve to the mean firing rates corresponding to cursor velocity directions (Ganguly and Carmena, 2009; Ganguly et al., 2011; Georgopoulos et al., 1988; Orsborn et al., 2014; Serruya et al., 2002). The MD of each neuron was then determined by taking the peak-to-peak amplitude of the cosine model. As predicted, we found that the MD was tightly coupled to the PV (Figure 4A) (correlation; Monkey C, R = 0.86, p < 1e-5; Monkey G, R = 0.63, p = 7e-3). Furthermore, the changes in MD were coupled to the changes in PV and was also able to change on a within-day timescale (Figures 4B, C) (correlation; Monkey J, R = 0.63, p = 0.0013; Monkey C, R = 0.73, p = 1e-4). Notably, regression lines very closely intersected the origin indicating that MD does not inherently increase due to task learning but could be due to modulation of PV. These results suggest that neurons are able to quickly increase the firing rates specifically in their preferred directions in response to perturbations in the control of the BMI. This increase in MD can be thought of as a “stretching” of the tuning curve or an amplitude gain that increases the variance of firing rate in the preferred direction without necessarily affecting the overall mean firing rate of the neuron.

**Figure 4:**
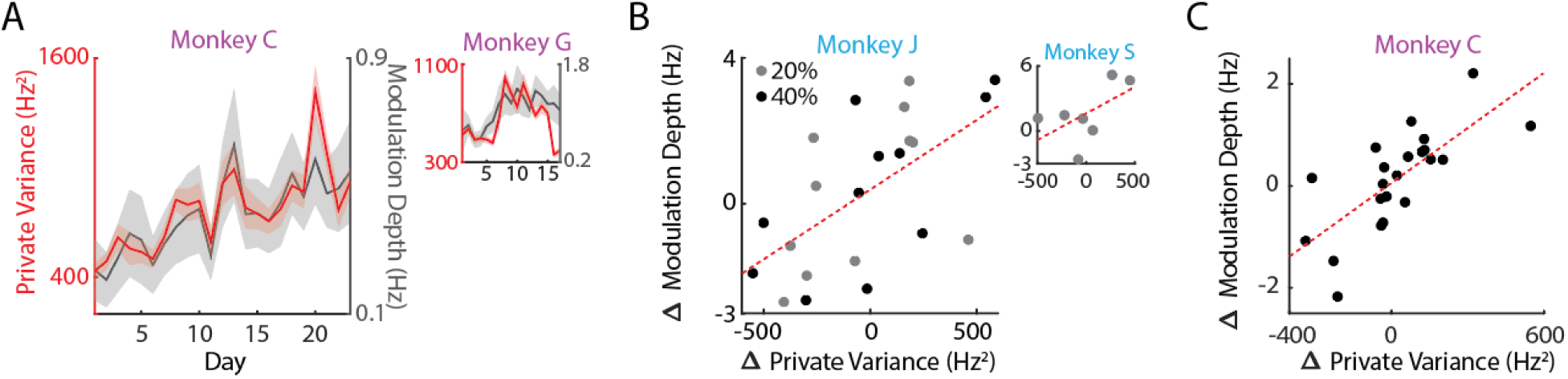
Modulation depth may facilitate PV changes. A. PV fluctuations are proportional to changes in MD (correlation; Monkey C, R = 0.86, p < 1e-5; Monkey G, R = 0.63, p = 7e-3). Lines indicate mean values of PV and MD; shaded regions show s.e.m. across targets for each day. While the changes in PV and MD were not significantly correlated for Monkey G in (C), the overall trends are proportional. B. Changes in PV were closely linked with changes in modulation depth (MD). Each dot represents one decoder change. Regression was conducted on combined data across the 24 days for Monkey J (correlation; R = 0.63, p = 0.0013). Correlation was not significant for Monkey S due to too few data but still shows same trends (correlation, R = 0.569, p = 0.18). C. Same as in (B) for Monkey C in Experiment 3 (correlation; R = 0.73, p = 1e-4). Results were not significant for Monkey G due to large fluctuations in channels used for recording from day-to-day.

## DISCUSSION

Previous work has shown BMI performance to improve over days alongside decoder re-adaptations (Oby et al., 2019; Orsborn et al., 2014). Here, we asked how neurons changed firing patterns over learning to allow for these performance improvements despite changing decoder weights. Our results support our hypothesis that motor cortex flexibly modulates neural variance when errors arise (due to decoder re-adaptation) without eroding preexisting coordinated patterns useful for generating goal-potent actions. Thus, at least in a 2-learner context, BMI learning involves leveraging exploratory patterns to adapt learned neural patterns rather than generating new ones.

Across all three experiments and five monkeys, the shared over total variance ratio (SOT) was remarkably stable across learning while private variance (PV) reliably fluctuated whenever a different decoder was introduced (Figure 3). Moreover, changes in PV were dependent on how different the decoders were from previously learned control spaces (Figure 3E). This was particularly noticeable in Experiment 1 where PV drastically changed within the same day while SOT remained unchanged within and across days. Notably, SOT did increase over longer timescales (i.e. Experiments 2 and 3) where perturbations were smaller. Experiment 1 used two separate decoders, with a new one introduced after achieving expert control with the first decoder. Furthermore, decoder perturbations happened each of the four days, each day ending with the previously learned decoder. The frequency and magnitude of these perturbations may have hindered consolidation effects observed in the other two experiments. Nonetheless, the SOT was stable within and across days and did not decrease due to repeated perturbations.

Also as predicted, we observed no changes in the low-dimensional neural subspace both within-day and across days (Supplemental Figure 2B-D). To quantify any changes, we calculated the “shared alignment” which represented the magnitude of the projection of one manifold onto another. In all three experiments, the alignment between any two adjacent blocks (or days for Experiments 2 and 3) were roughly 0.8∼0.9 indicating similar subspaces on short timescales. These alignments were also similar across days suggesting minimal rotation of the manifold even over the time-course of learning.

Our results also suggest that modulation depth (MD) may play a critical role in facilitating fast changes in neural variance. While past studies have repeatedly shown changes in MD over learning, it has been unclear what the exact role of this may be. We found that the MD was tightly correlated with PV. More specifically, changes in PV were closely tracked by directly proportional changes in MD (Figure 4). Importantly, fluctuations in both were fast (compared to neural consolidation rates) in both increasing and decreasing directions (Figures 2, 4, Supplemental Figure 1). Intuitively, increasing MD increases the dynamic range of neural firing rates, allowing for higher total neural variance. In contrast to past studies showing overall changes in MD over learning (Ganguly and Carmena, 2009; Ganguly et al., 2011; Orsborn et al., 2014), these fast changes in modulation depth indicate that the ability to fluctuate firing rates may be an inherent property of motor cortex rather than a learned behavior.

Lastly, our findings reflecting flexible changes without greatly affecting established coordinated patterns is compatible with that of natural motor adaptation. Our results suggest that neuroprosthetic learning and motor adaptation may have similar underlying mechanisms. Studies have shown the importance of subcortical structures and the cerebellum in adaption to motor perturbations and BMI tasks (Koralek et al., 2012; Nowak et al., 2007; Shadmehr and Krakauer, 2008). Futhermore, past studies provide evidence on the role of the basal ganglia for refining cortical patterns during BMI on shorter and longer timescales (Athalye et al., 2018, 2020; Costa et al., 2004; Ölveczky et al., 2011). In a motor adaptation task then, it is possible these structures are turning the “variability knob” in motor cortex. If we consider the production of motor actions as a combination of consolidated activity from the motor system with some exploratory variability (Fee and Goldberg, 2011; Neuringer et al., 2000), then our results suggest that PV may be a neural correlate of the latter. Furthermore, changes in neural variance might reflect an increased gain in exploratory variability without directly affecting learned motor behaviors. In such a scenario, neuroprosthetic learning – particularly with closed-loop decoder adaptation (CLDA) – may be encouraging changes from subcortical structures we see manifested as changes in motor cortex.

Our results are complementary with those from previous studies (Athalye et al., 2017; Golub et al., 2018; Oby et al., 2019; Sadtler et al., 2014). It has been shown that intrinsic manifolds are relatively stable and thus are less able to adapt to decoders that are explicitly out of that manifold (Sadtler et al., 2014). While learning out-of-manifold decoders is difficult within-day, it has also been shown that these shared spaces can rotate over many days of training (Athalye et al., 2017; Oby et al., 2019). In this study, we show how neurons respond to changes in decoder weights on both timescales before task performance is saturated. Our results regarding stable shared spaces corroborate these past studies but we highlight the importance of the neural variance (and specifically the PV) during learning. While we echo past results that neural activity preferentailly stays in a constant neural subspace during learning, we found that neurons are also able to quickly increase their private variance when perturbations are introduced. This simultaneous modulation of neural variance whilst maintaining a robust manifold suggests a neural learning strategy that is resilient to perturbations.

## METHODS

### Surgical Procedures and Electrophysiology

All procedures for each of the three experiments were conducted in compliance with the NIH *Guide for the Care and Use of Laboratory Animals* and were approved by the University of California, Berkeley Institutional Animal Care and Use Committee.

#### Experiment 1

One adult male rhesus macaque was chronically implanted with 64-channel Teflon-coated tungsten microelectrode arrays. Implants were in the arm regions of primary motor cortex (M1) and dorsal premotor cortex (PMd) in the left hemisphere. There was an additional implant in the arm area of M1 in the right hemisphere, totaling three implants. Unit activity was recorded using a MAP system (Plexon Inc., Dallas, TX) and sorted with an online spike-sorting application (Sort Client; Plexon). Units stable across days were used for decoding (Ganguly and Carmena, 2009).

#### Experiment 2

Two adult male rhesus macaques were chronically implanted with 128-channel microwire electrode arrays. Implants were bilateral, targeting the arm areas of M1. Neural activity was recorded using the same MAP system as in Experiment 1. Multi-unit activity was sorted using Sort Client for Monkey S and channel-level activity was used for Monkey J (Orsborn et al., 2014).

#### Experiment 3

Two adult male rhesus macaques were chronically implanted with microwire electrode arrays with 128 channels in each hemisphere, targeting arm areas of M1 and PMd in Monkey G and primary sensory cortex (S1) and M1 in Monkey C. Neural activity was filtered and recorded using an OmniPlex system (Plexon Inc., Dallas, TX). Single and multi-unit activity was sorted with a template-matching algorithm (PlexControl) (Moorman et al., 2017).

### BMI Task

In all three of the experiments we analyzed, monkeys performed a center-out BMI task. Subjects were instructed to enter and hold in the center target to initiate a trial. Upon hold completion, one of eight peripheral targets would appear. Subjects then moved the cursor towards the target after this go cue. Trials were successful if the cursor entered and stayed in the peripheral target for a short hold time (∼250 ms) for each experiment. Decoders were trained differently in the three experiments (outlined below), but in all cases, subjects were familiar with the task structure before attempting control with BMI. Therefore, subjects’ performance improvement can be attributed to BMI learning rather than task learning. Once decoder weights were held fixed each day, task control was achieved by modulating neural activity, which was recorded and fed through the decoder to produce cursor movements. In order to observe neural correlates of coordinated control, we analyzed movement portions (i.e. leaving center target to entering peripheral target) of successful trials for each experiment (red box, Figure 1B).

### Experiment 1

In Experiment 1, two macaques were instructed to complete a BMI center-out task as described above. After achieving proficient control with an “old” decoder, a new biomimetic decoder was introduced. This decoder was trained on Day 1 (Figure 1 C). While many weights were similar between the two decoders, many of the weights were significantly different (Ganguly and Carmena, 2009). Each day, both decoders were used for BMI control in a series of two blocks. To track proficiency with each decoder, the percent of correct trials was calculated for each block for each day. For each decoder, a linear filter (Wiener filter) was used to fit neural activity to kinematic activity in the elbow and shoulder joints during a manual reaching task using a BKIN KINARM exoskeleton robot. Decoder outputs were then translated into Cartesian space by using a Jacobian matrix. For more details, we redirect the reader to Ganguly and Carmena, 2009.

#### Wiener Filter

The Wiener filter model assumes that the cursor velocity is a linear combination of neural activity across small differences in time. In other words, the output of the decoder at time *t* is determined by a weighted sum of the neural activity from *t* − *k* for some set of *k* representing the time lags. More specifically:

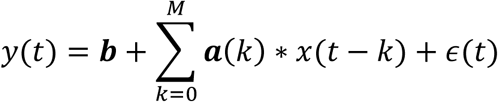

where *ϵ* refers to any residual errors and *M* represents the number of time lags used (in Experiment 1, this was 10). The parameters ***a*** and ***b*** were determined to optimally fit the model (Ganguly and Carmena, 2009; Wessberg and Nicolelis, 2004).

### Experiment 2

The second experiment we analyzed was originally conducted by Orsborn et al in 2014. To summarize, prior to BMI control, both subjects were trained to conduct the manual control version of the center-out task using the KINARM. An initial Kalman filter (KF, outlined below) decoder was fit on day 1 on these manual reaches. Closed-loop decoder adaptation (CLDA) was then used to better fit decoder weights to the neural intentions (Orsborn et al., 2012). To briefly summarize, CLDA updates decoder weights by inferring intentions of the subject based on task goals. Decoder weights were updated through a weighted average of old and new decoder weights. CLDA was run at the beginning of day 1 to achieve some level of control (∼20% accuracy). Over days, decoder weights were occasionally refit with CLDA, sometimes in conjunction with neuron swaps to recover performance. After 12 days, the procedure was repeated for Monkey J with initial performance elevated to 40% accuracy. For further details regarding CLDA, we redirect readers to the methods outlined in Orsborn et al., 2012.

### Experiment 3

In Experiment 3, two monkeys (Monkeys C and G) controlled a four degree-of-freedom (DOF) kinematic chain in a center-out task. Subjects were first trained to perform a cursor center-out task using manual control with a KINARM exoskeleton as in previous studies, then were transitioned to BMI control of a 2D cursor before learning to perform the full 4 DOF task with BMI control. Movements of the kinematic chain were constrained to two dimensions thereby creating a kinematically redundant system. A KF decoder fit neural activity to four individual joint angle velocities. Visual feedback of the kinematic chain moving from target to target was used to seed the KF decoder and CLDA was used to update neural intentions each day. Unlike Experiment 2, CLDA was run until the subjects’ performance saturated each day (∼85% accuracy). Since the percent correct was saturated and stable across days, task improvement was tracked by decreasing reach times. We refer readers to Moorman et al., 2017 for more experimental details.

#### Kalman Filter

Here, we briefly describe the Kalman filter model for BMI decoding used in Experiments 2 and For more details, we refer the reader to Orsborn et al., 2014 and Moorman et al., 2017. In general, the state-space (e.g. kinematic space) is represented by *x*_*t*_ = [*p*_*t*_ *v*_*t*_ 1] where *p*_*t*_ represents the positions of the effector and *v*_*t*_ represents the velocities of the effector. In Experiment 2, the state-space contains the horizontal and vertical positions and velocities. In Experiment 3, the state-space refers to the joint angles and angular velocities of the 4 DOF kinematic chain. The Kalman Filter model assumes that *x*_*t*_ varies as *x*_*t*+1_ = **A***x*_*t*_ + *w*_*t*_ where *w*_*t*_ is a noise term (*w*_*t*_ ∼ *N*(0, **W**)). Aadvances the state using the current positions and velocities and *W* is set up so as to allow the joint velocities to evolve independently:

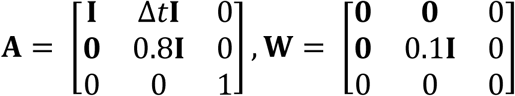

where **I, 0** are identity and 0 matrices, respectively, and Δ*t* is 100 ms. Note that in these models, **A** contains a scaling factor (0.8) in order to decay the velocities over time.

The observations *y*_*t*_ represent the neural activity in the past 100 ms. The KF model of neural firing assumes that *y*_*t*_ = **C***x*_*t*_ +***q***_*t*_, ***q***_*t*_ ∼ *N*(0, ***Q***). The KF estimates *x*_*t*_, the state variable, from the previous tobservations {*y*_0_, …, *y*_*t*_ } and produces the state estimate *x*_*t*_ and confidence **P**_*t*_. The KF uses the last estimate *x*_*t* − 1_ and advances that belief to produce *x*_*t*| *t*−1_ =*Ax*_*t*−1_, and then updates the prediction when a new observation becomes available: *x*_*t*_ *= x*_*t*| *t*−1_ +*K*_*t*_ (*y*_*t*_ –*Cx*_*t* | *t*−1_). The Kalman gain **K**_*t*_ is then determined by the model parameters {**A, C, W, Q**} and P_*t*_. Since our model parameters are fixed, **K**_*t*_ converges to a steady-state **K**, and we estimate the kinematics to follow the simpler equation:

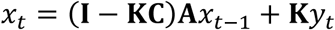

### Factor Analysis

Factor analysis (FA) was used in this study to approximate low-dimensional manifolds; here we provide an outline of the method (Athalye et al., 2017). FA decomposes the neural activity into three components: the mean firing rates (*μ*) of each neuron, the population-level coordinated firing activity (*x*^*shared*^), and the remaining component corresponding to each neuron’s uncorrelated activity to the rest of the population (*x*^*private*^). That is,

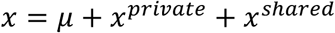

These latter two components have covariance matrices Σ^*private*^ and Σ^*shared*^ with dimensions *N* × *N*, where *N* is the number of neurons. The combination of these two parts yields the total variance,

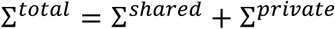

The dimensionality was estimated by using cross-validated log-likelihood to determine the number of factors that would best describe held-out data such that ∼90% of the shared variance could be captured (Athalye et al., 2017).

Importantly, only units stable across all days for each experiment were used for these analyses. Since we were interested in how the same ensemble of units adapted over learning, including all units used for each day would contaminate across-day results.

### Private Variance

We define private variance (PV) as an *N* × *N* covariance matrix where the diagonals represent independent variances for each of the *N* neurons. In Experiment 1, the PV was calculated for each target and each block. Average PV values were found by averaging across the 8 targets for each block. Similarly, PV was calculated for each day (and one day after) a decoder was refit in Experiment 2. This gave us PV values before and after refitting and a t-test was used to determine differences in PV. Furthermore, to track how PV changed over days with stable decoders, PV was calculated for days without decoder changes as well. In Experiment 3, since CLDA was run each day of the experiment and percent accuracy on Day 1 was already as good as the last day (i.e. proficient control), we instead examined the correlation of the decoder for each day with the decoder on the first day as a metric for the difference between decoders. We then plotted this correlation against the average private variance over all neurons for that day (Figure 3E). Note that this correlation was shown for stable units in Figure 3E; Supplemental Figure 3A shows this correlation calculated using stable channels for Monkey G, which include channels that were used in the BMI task but were either physiologically different across recording days or contained no physiological signal.

### Shared / Total Variance

We defined the shared/total variance ratio (SOT) as

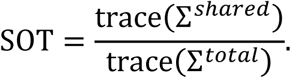

Intuitively, this allowed us to measure the proportion of neural variance that was coordinated. In order to track how neurons in the population changed coordination patterns over learning, we only considered units that were stable across the entirety of the experiments for these analyses, determined by waveform and inter-spike interval statistics. In all experiments, the SOT was calculated for each block/day for all stable units. In Experiment 3, Monkey G had several “unstable” units throughout the experiment which were not considered in these analyses. In addition, a variety of noisy channels were included as decoder inputs. Figures 3 and 4 show results with stable units for Monkey G, however the results still hold using all stable channels (Supplemental Figures 4-5).

### Shared Space Alignment

In order to determine how similar manifolds were between experimental blocks (or days), we calculated the “alignment” between shared spaces (Athalye et al., 2017). Geometrically, it shows how much of one subspace projects onto a second as a fraction ranging from 0 to 1. Moreover, a perfectly aligned subspace results in an alignment value of 1 while orthogonal subspaces have an alignment of 0. To calculate the shared space alignment, we find the projection of the shared variance of block (or day) *A* onto the shared space of block (or day) *B*.

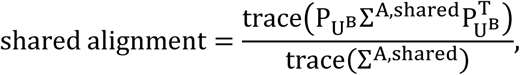

where 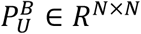 is an projection matrix onto the shared subspace of block *B* (col(*U*^*B*^)), and Σ^*A*,shared^ is the shared variance of block *A*.

To measure how similar shared spaces were between decoder changes, we first found the shared alignment between pairwise days. In Experiment 1, comparisons were then made between all decoder block changes and all other pairwise comparisons. Similarly, to test if shared spaces were aligned on short and longer timescales in Experiments 2 and 3, shared alignments between adjacent days were compared against all other pairwise combinations of days. For each experiment, a Kolmogorov-Smirnov test was used to determine if the distributions of shared alignment values were significantly different.

### Preferred Direction and Modulation Depth

We also looked at the correlation of modulation depth and private variance. Cursor kinematics were separated into eight 45-degree bins and corresponding firing rates were used to determine the preferred tuning direction (PD) for each neuron by fitting the firing rate *f* to a cosine function with mean firing rate *μ*:

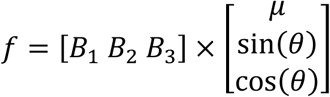

where *θ* corresponds to cursor movement angle and **B** are coefficients estimated through linear regression. PD was calculated as PD = arctan(*B*_2_ /*B*_3_), resolved to the correct quadrant. Modulation depth (MD) was then defined as the peak-to-peak amplitude of this curve. Intuitively, it shows how specific neurons fire in their PD and is calculated as

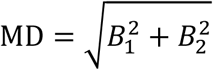

To track how MD and PV changed together over learning, we correlated the changes in MD (i.e. derivative) with the changes in PV across days. As before, Figure 4A (inset) shows the results when considering only stable units for Monkey G to remain consistent with the other animals. However, the results are consistent if all stable channels are used as well (Supplemental Figure 3B).

### Statistical Analyses and Testing

We used t-tests to reject null hypotheses with a confidence of 5% and used linear regressions and correlations to fit trends. Kolmogorov-Smirnov tests were used to test whether distributions were the same. Wilcoxon rank-sum tests were used to compare private variance distribution differences. Where reported, error bars represent standard error of the mean over 8 targets. All analyses were done in MATLAB by writing custom scripts.

