## Supplemental Figures for "Flexible Modulation of Neural Variance Facilitates Neuroprosthetic Skill Learning"

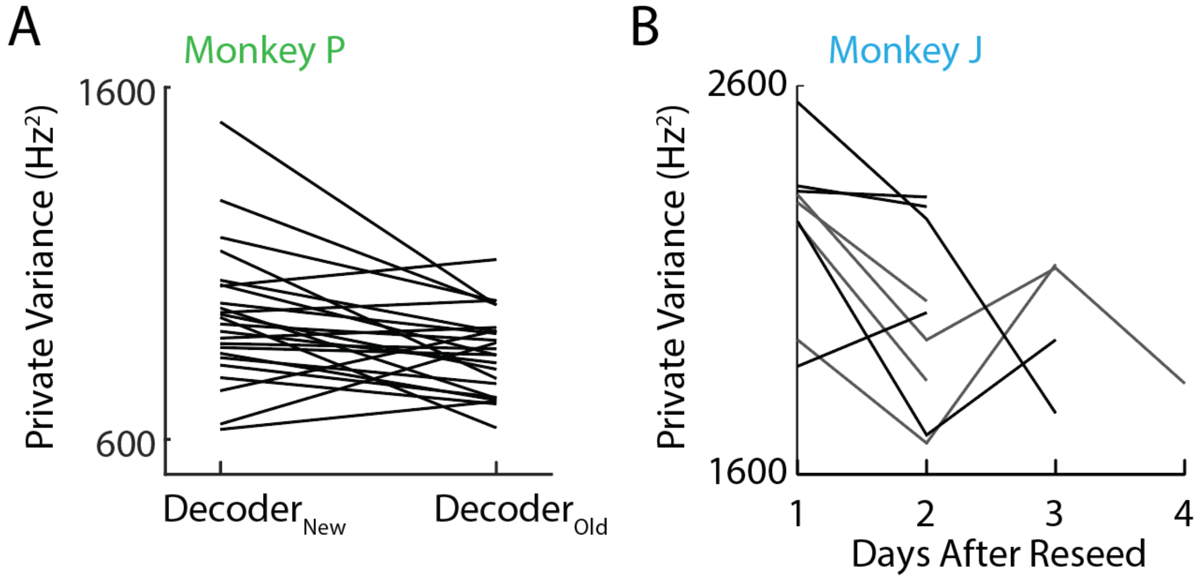


**Supplemental Figure 1: Private variance decreases over learning with a fixed decoder.**

A. Private variance decreased for Monkey P when the task switched from using the unfamiliar decoder to the familiar decoder. Each line indicates changes in PV for a single target for each day (t-test, p = 0.008).

B. When decoder weights were constant in Experiment 2, PV decreased (Wilcoxon rank-sum test, p = 0.0315).

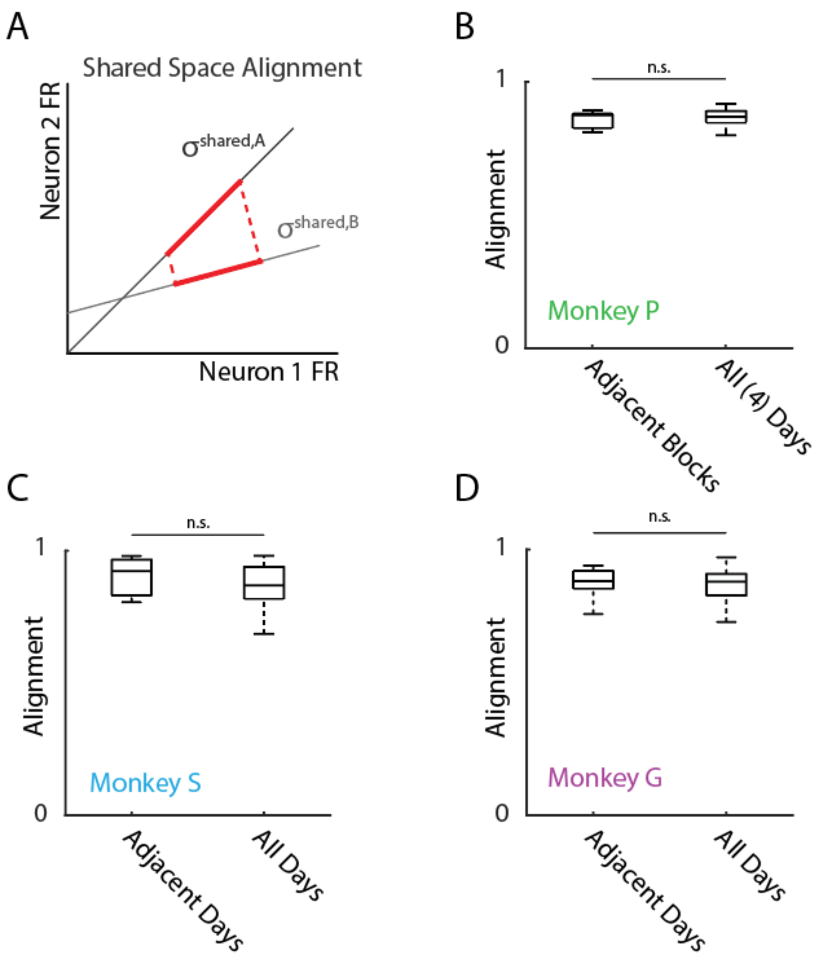


**Supplemental Figure 2: Shared spaces are resilient to perturbations**

A. The similarity (alignment) between shared spaces was calculated by projecting the shared space of one experimental block onto another. The shared space alignment was defined as the amount of variance captured by this projection.

B. Alignment was calculated between adjacent blocks in Experiment 1 as well as all pairwise comparisons among all blocks across the four days. The shared alignment was high between decoder swaps indicating a stable space immediately following a decoder change. This shared alignment was not significantly different between adjacent days and over the time course of four days ($p > 0.05$, Kolmogorov-Smirnov test).

C. Same as B except over days rather than individual blocks for Experiment 2. Shared alignment was not significantly different between short and long time periods ($p > 0.05$, Kolmogorov-Smirnov test).

D. Same as C but for Experiment 3 ($p > 0.05$).

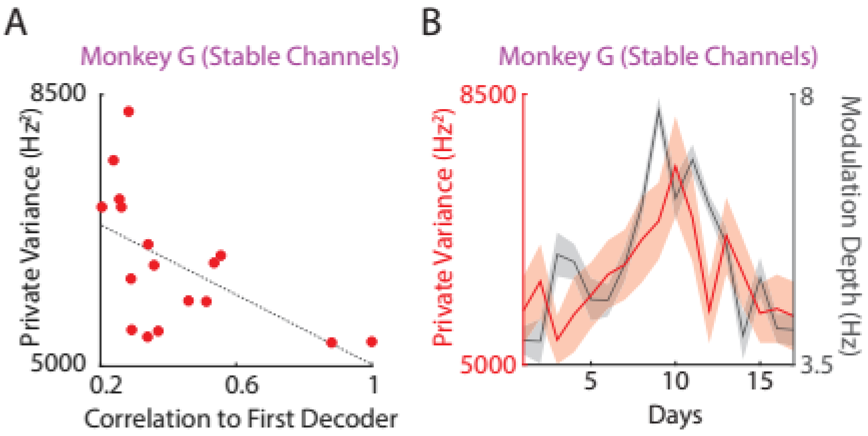


**Supplemental Figure 3: Private variance modulation in stable channels for Monkey G**

A. Same as Figure 3E but with stable channels in Monkey G (correlation; R = -0.57, p = 0.0169).

B. Same as Figure 4A inset but with stable channels in Monkey G (correlation; R = 0.61, p = 0.0088).

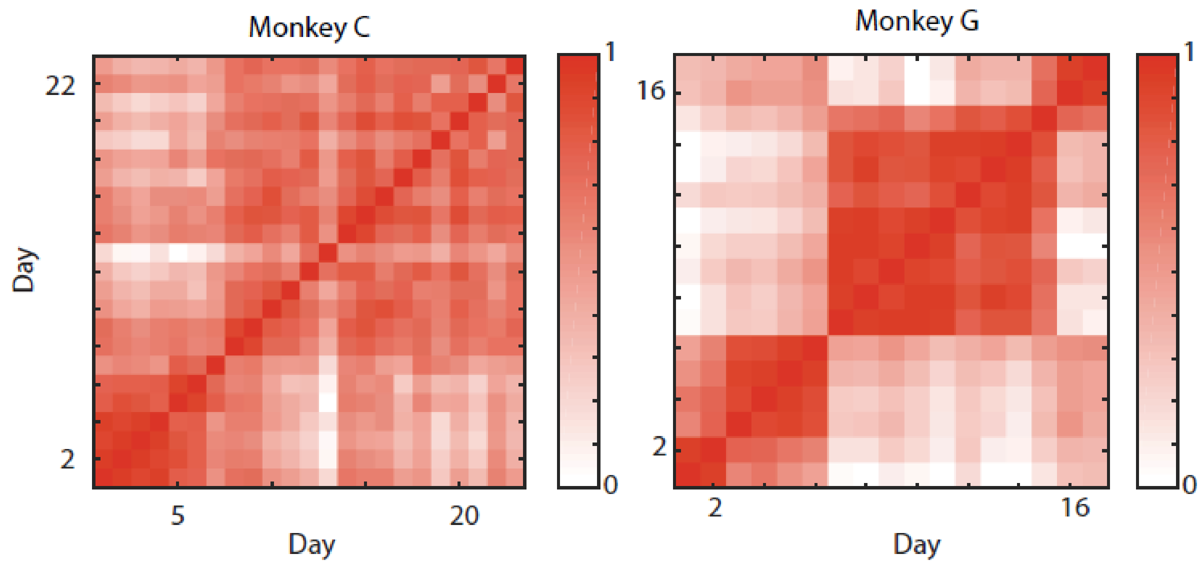


**Supplemental Figure 4: Decoder correlations**

Decoder observation model weights were correlated between pairwise days for Monkey C (left) and Monkey G (right). Only stable units were used in this plot for Monkey G. Decoder correlations across all stable channels for Monkey G are shown in Supplemental Figure 5.

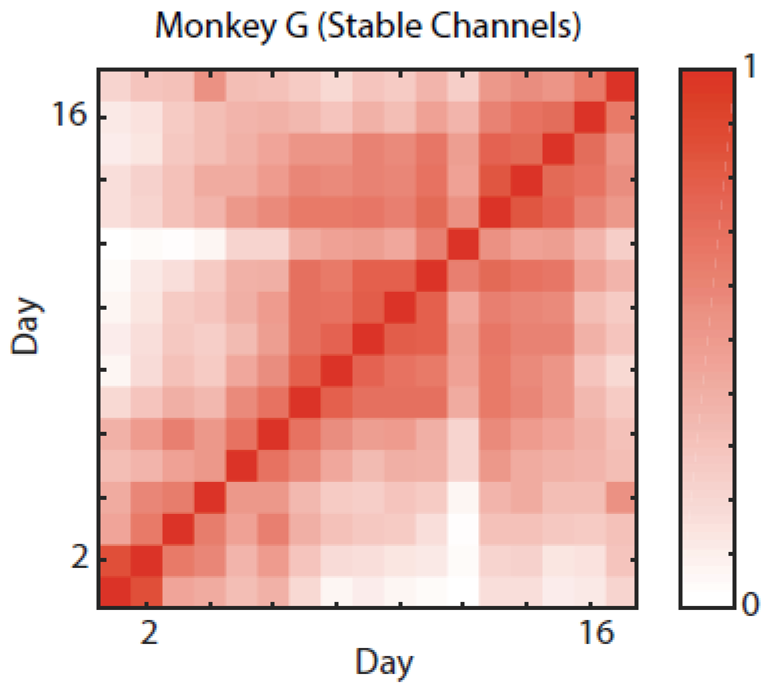


**Supplemental Figure 5: Decoder correlations for stable channels in Monkey G**

Same as Supplemental Figure 4, but for all stably recorded channels in Monkey G.
